# Cardiac Microlesions Form During Severe Bacteremic *Enterococcus faecalis* Infection

**DOI:** 10.1101/2020.01.14.906669

**Authors:** Armand O. Brown, Kavindra V. Singh, Melissa R. Cruz, Karan Gautam Kaval, Liezl E. Francisco, Barbara E. Murray, Danielle A. Garsin

**Affiliations:** Department of Microbiology and Molecular Genetics, The University of Texas Health Science Center at Houston, Houston TX, USA; Division of Infectious Diseases, Department of Internal Medicine, The University of Texas Health Science Center at Houston, Houston, TX, USA; Department of Biochemistry and Structural Biology, The University of Texas Health Science Center at San Antonio, San Antonio, TX, USA

**Keywords:** Enterococcus faecalis, cardiac, microlesions, sepsis, bacteremia, infection, virulence, thioredoxin, disulfide bond, necroptosis

## Abstract

*Enterococcus faecalis* is a significant cause of hospital-acquired bacteremia. Herein, the discovery is reported that cardiac microlesions form during severe bacteremic *E. faecalis* infection in mice. The cardiac microlesions were identical in appearance to those formed by *Streptococcus pneumoniae* during invasive pneumococcal disease (IPD). However, *E. faecalis* does not encode the virulence determinants implicated in pneumococcal microlesion formation. Rather, disulfide bond forming protein DsbA was found to be required for *E. faecalis* virulence in a *C. elegans* model and was necessary for efficient cardiac microlesion formation. Furthermore, *E. faecalis* promoted cardiomyocyte apoptotic and necroptotic cell death at sites of microlesion formation. Additionally, loss of DsbA caused an increase in pro-inflammatory cytokines unlike the wild-type strain, which suppressed the immune response. In conclusion, we establish that *E. faecalis* is capable of forming cardiac microlesions and identify features of both the bacterium and the host response that are mechanistically involved.

**SUMMARY:** This work presents the observation of cardiac microlesion formation during severe blood stream infection with *Enterococcus faecalis* in mice. Moreover, we identify the contribution of a novel enterococcal virulence determinant in modulating microlesion formation and the host immune response.

## Introduction

*Enterococcus faecalis* is a Gram-positive bacterium that normally co-exists with the host as a harmless commensal. However, it is capable of causing a variety of infections such as infective endocarditis (IE), urinary tract infection (UTI), bacteremia, peritonitis, prosthetic joint infection, and endophthalmitis. A common complication associated with enterococcal bacteremia is endocarditis, an infection that has a 1-year mortality rate of approximately 29% despite antibiotic therapy [1]. Enterococci are the third most common cause of endocarditis in North America [2] and, in patients with this infection, heart failure has been observed to occur with increased frequency [3–9]. Endocarditis only involves infection of the surfaces of the heart - the formation of biofilms on heart valves and inner chambers that develop into complex masses called vegetations [10]. *E. faecalis* invasion of the deeper myocardial tissue to cause cardiac microlesions has not previously been described.

Interestingly, another Gram-positive pathogen, *Streptococcus pneumoniae*, was recently shown to invade the heart tissue during invasive pneumococcal disease (IPD) [11]. Importantly, pneumococcal microlesions were determined to be associated with altered cardiac electrophysiology, heart failure and scar formation during convalescence, which is thought to predispose individuals to major adverse cardiac events (MACE) [11, 12]. Furthermore, cardiac microlesions form following vascular endothelial cell invasion into the myocardial tissue through the interactions of pneumococcal Choline binding protein A (CbpA) and pneumococcal teichoic acid associated phosphorylcholine (ChoP) with host laminin receptor (LR) and platelet activating factor receptor (PAFr), respectively. Cardiomyocyte death is mediated by exposure to the pneumococcal secreted cholesterol-dependent pore forming toxin, pneumolysin [11, 12], and hydrogen peroxide [13, 14]. Since the original observation of pneumococcal microlesions in 2014, similar observations were seen during blood stream infections with *Francisella tularensis* [15], and *Mycobacterium avium* [16], however the mechanisms involved were not characterized.

In this report, we show that cardiac microlesions also form during invasive *E. faecalis* infection. The observed cardiac microlesions were identical in appearance and size to those formed by *S. pneumoniae*, were diffuse across the myocardium, and characterized by an absence of immune cell infiltration and cell death by apoptosis and necroptosis. Furthermore, an enterococcal determinant required for microlesion formation, DsbA, was identified. DsbA was additionally shown to enhance cell death and promote suppression of the immune response during *E. faecalis* infection of a cardiomyocyte cell line.

## Materials and Methods

### *C. elegans* survival assay

*E. faecalis* strains were tested in *C. elegans* for survival as previously described [17]. The data is representative of 3 independent experiments. Nematode survival was plotted with the log-rank (Mantel-Cox) method using Prism 7 statistical software (GraphPad).

### Bacterial strains and growth analysis

All *E. faecalis* strains used in this project are listed in Table S1. For growth analysis, overnight cultures were diluted to an OD_600nm_ of 0.05 in brain heart infusion (BHI) medium (Difco) or serum [18, 19] in a volume of 200µl / well in a 96-well plate. The 96-well plate containing the diluted cultures were placed in a prewarmed 37°C BioTek Cytation 5 plate reader, and OD_600nm_ measurements were taken every 10 minutes for 16 hours.

### Mouse peritonitis and cardiac microlesion model

The mouse peritonitis model was performed as described previously [20]. The method for identifying and evaluating cardiac microlesion formation was conducted as described previously [11]. Briefly, female, ~4 – 6-week old (~25g), ICR outbred mice were obtained from Envigo (Indianapolis, Indiana) and infected intraperitoneally with 3.8 - 4.2 x10^8^ CFUs of *E. faecalis*. Mouse hearts were harvested 24-48 hours post-infection at the point when the animals showed signs of severe sepsis indicated by ruffled fur, labored breathing and little movement. The hearts were formalin fixed and sent to The University of Texas Health Science Center Histopathology Laboratory for paraffin embedding, sectioning, and hematoxylin and eosin H&E) staining. Paraffin embedded mouse hearts were first bisected and sectioned longitudinally. 20 sections were cut per block, and every 5^th^ section H&E stained in order to measure the number and relative distribution of the microlesions. Cardiac microlesions were averaged across sections, and the differences between mouse groups compared using *Student’s* two-tailed t-test. Experiments were performed in accordance with protocols approved by the Animal Welfare Committee of the University of Texas Health Science Center at Houston AWC-19-0058.

### Invasion assay

SVEC4-10 Endothelial Cells (ATCC) were seeded in 24 well plates at a cell density of 1.5×10^5^ cells/ml. After 24 hours, the cells were washed three times with PBS and infected with *E. faecalis* at a MOI of 10 for 4 hours in antibiotic free Dulbecco’s Modified Eagle’s Medium (ATCC). The cells were then washed with Dulbecco’s (1X) PBS three times, treated with trypsin, diluted in PBS, and plated for quantification. To measure intracellular numbers of *E. faecalis*, cells that were washed three times with Dulbecco’s (1X) PBS 4 h.p.i., then treated with DMEM (ATCC) supplemented with 15 µg/ml vancomycin and 150 µg/ml gentamicin. 24 hours later, the wells were washed three times with Dulbecco’s (1X) PBS, lysed using 0.1% Triton-X-100, and diluted and plated for intracellular quantification. The ratio of invasion to attachment was used to calculate the percent of invasion. The experiments were conducted on 3 independent occasions and were performed using triplicate wells for each strain. Attachment and invasion values were averaged from each experiment, and statistically analyzed using *Student’s* two-tailed t-test.

### Cytotoxicity and inflammatory cytokine panel microtiter plate reader assay

HL-1 cardiomyocytes were obtained from Sigma-Aldrich and SVEC4-10 Endothelial Cells were obtained from ATCC. HL-1 cardiomyocytes (Sigma-Aldrich) were seeded into 24-well plates at a concentration of 5×10^5^ cells/ml. 24 hours later, the cells were washed three times with Dulbecco’s (1X) PBS, and infected with *E. faecalis* at a MOI of 10 for 4 hours in antibiotic free Claycomb medium. Supernatant was collected, and cytotoxicity assayed using CytoTox-ONE™ Homogeneous Membrane Integrity Assay Kit (Promega). For HL-1 infected cardiomyocytes, inflammatory chemokines and cytokines were additionally analyzed using a Mouse Cytokine ELISA Plate Array III Colorimetric Assay (Signosis). The data represent an average of at least 3 independent experiments for each strain and were analyzed using *Student’s* two-tailed t-test.

### Immunofluorescence microscopy

Immunofluorescence microscopy was conducted as described previously [11]. Briefly, paraffin embedded sections were treated with 1:100 rabbit Anti-mouse MLKL (phospho S345) antibody [EPR9515(2)] in order to detect phosphorylated MLKL (Abcam), and 1:100 rat anti-mouse MLKL antibody (Millipore-Sigma) for MLKL detection followed by treatment with 1:1000 chicken anti-rabbit IgG (H+L) cross-adsorbed secondary Alexa Fluor 594 antibody (ThermoFisher), and 1:1000 goat anti-rat IgG (H+L) secondary FITC antibody (ThermoFisher) respectively. A rabbit anti-*Enterococcus faecalis* antibody (Abcam) was used at a concentration of 1:1000 to detect *E. faecalis* followed by treatment with 1:1000 goat anti-rabbit IgG (H&L), FITC conjugated antibody (Invitrogen). A rat anti-PARP (MAB600) antibody (R&DSystems) was used at a concentration of 1:100 followed by treatment with 1:1000 goat anti-rat IgG (H+L) cross-adsorbed secondary Alexa Fluor 594 antibody (ThermoFisher) to detect PARP. All stained sections were counterstained with 1:1000 DAPI (ThermoFisher) and mounted with Hardset™ Antifade Mounting Medium. Visualization of fluorescence was obtained using standard excitation and emission wavelengths for Alexa 594, FITC, and DAPI on an Olympus FV3000RS high-speed, high-sensitivity, inverted laser scanning confocal microscope.

### Generation of rabbit anti-DsbA antisera

His-tagged recombinant DsbA was overexpressed in Rosetta-2(DE3) *E. coli* (Sigma-Aldrich) and isolated by affinity purification via HisPur Nickel-NTA Resin (ThermoFisher) packed columns followed by Fast Protein Liquid Chromatography (FPLC) to obtain highly pure samples of recombinant DsbA. Cocalico Biologicals (Reamstown, PA) generated anti-DsbA atisera by immunizing rabbits with the recombinant protein.

### *E. faecalis* whole cell lysate and Western blot

A 5 ml culture of *E. faecalis* grown in BHI medium at 37°C for 15-16 hours followed by a 1:10 dilution in 5 ml of fresh BHI. After 3.5 hours to achieve exponential growth, the culture was centrifuged at 6,010 × G and the supernatant removed. The pellet was resuspended in 1ml of buffer (10 mM Tris/HCl pH 7.5 and 200 mM NaCl) containing cOmplete^™^, EDTA-free Protease Inhibitor Cocktail (Roche). Following sonication of the pellet on ice, the sample was centrifuged at 10,000 × G for 5 minutes and supernatant collected for electrophoresis. Total protein in the whole cell lysate was measured using the *DC* Protein Assay (BioRad). Following the loading of 10µg of cell lysate, electrophoresis and transfer of protein to a PVDF membrane, the membrane was incubated overnight at 4°C in a 5% milk solution containing a 1:1000 dilution of rabbit anti-DsbA antisera followed by incubation with a Goat anti-rabbit IgG HRP (ThermoFisher) secondary antibody at a dilution of 1:10,000 at room temperature for 30 minutes prior to development with a chemiluminescent kit (ThermoFisher).

## Results

### Cardiac microlesions form during severe bacteremic *E. faecalis* infection

In previous work, we implicated a previously uncharacterized *E. faecalis* thioredoxin known as DsbA (disulfide bond forming protein A) as a requirement for the stability of EntV, a secreted bacteriocin that inhibits hyphal morphogenesis of the human fungal pathogen *Candida albicans* [21, 22]. To test if these proteins had any effects on the fitness and pathogenicity of *E. faecalis*, we infected *Caenorhabditis elegans* with Δ*dsbA* and Δ*entV E. faecalis* mutants and measured survival. As shown in Figure 1A, loss of *dsbA* significantly attenuated mortality in infected nematodes. Note that the Δ*dsbA* mutant did not manifest different growth kinetics when grown in culture medium compared to wild-type, ruling out a general growth defect (Figure S1A). Infection with the complement, Δ*dsbA::dsbA*, partially rescued the Δ*dsbA* phenotype. The partial complementation was attributed to the DsbA protein being produced at lower levels in the complemented strain (Figure 1B and S2). No significant difference in nematode survival was observed following infection with either the Δ*entV* mutant or its complement (Figure 1A). Considering that EntV is the only known substrate of *E. faecalis* DsbA, the results suggest that DsbA has additional targets that affect pathogenicity.

**Figure 1.**
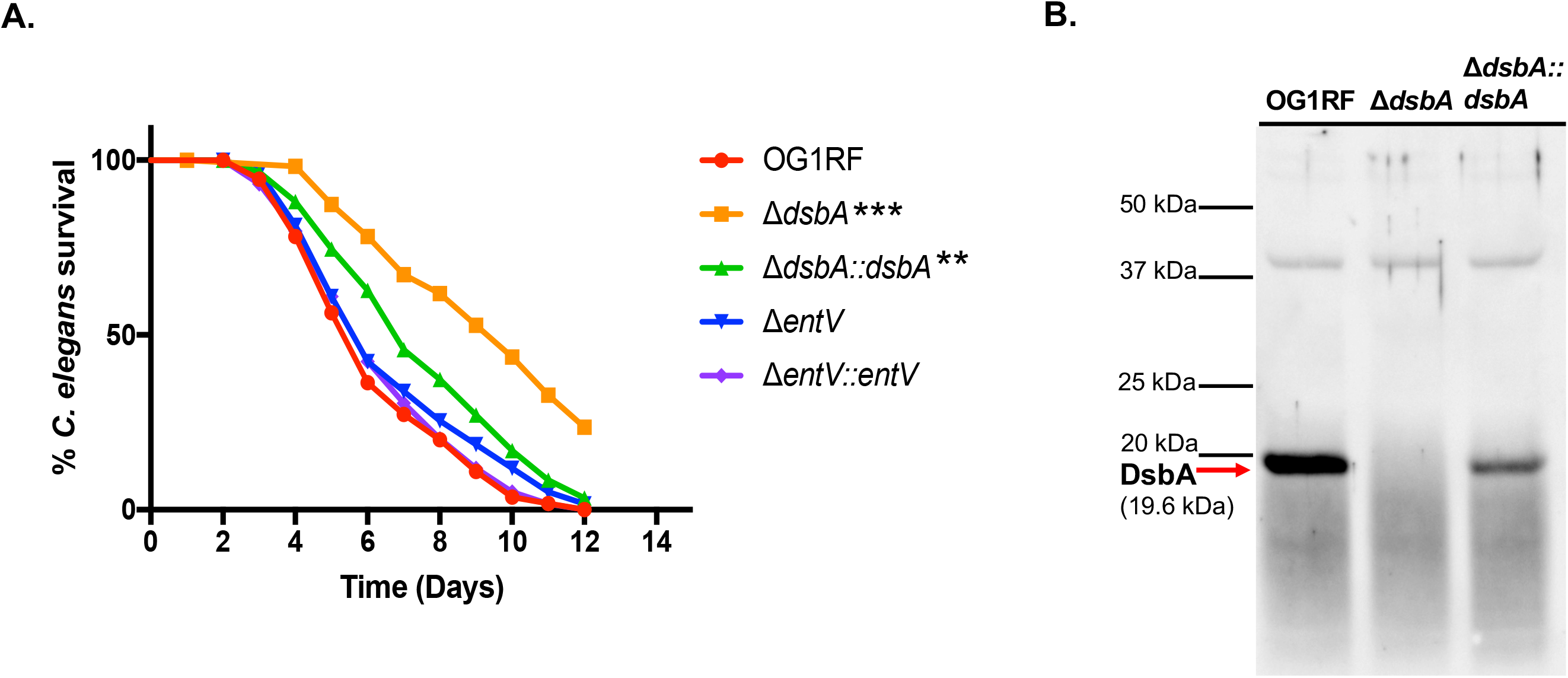
*dsbA* deficient *E. faecalis* show reduced virulence in a *C. elegans* survival assay. **(A)** L4 stage *C. elegans* were infected with *E. faecalis*. This figure is representative of 3 independent experiments with 30 worms/group. ****Asterisks indicate a *P* < 0.0001, ** a P < 0.01 when compared to OG1RF following pairwise analysis by log-rank (Mantel-Cox). **(B)** Western blot performed by SDS-PAGE separation of *E. faecalis* whole cell lysates followed by hybridization of DsbA using rabbit anti-DsbA antisera. Equal loading is ensured by visualization of a non-specific band at 40 kDa. Image is representative of 3 independent Western blot experiments.

Based on these results, we wanted to know if loss of DsbA also affected virulence in a vertebrate animal model of *E. faecalis* infection. Therefore, the mutants were tested in a mouse model in which animals were injected intra-peritoneally with *E. faecalis* resulting in generalized bacteremia leading to mortality. Significant survival differences between the mice infected with the different strains were subtle, but significant attenuation by the *dsbA* mutant was observed when data from three separate experiments were combined to generate an n close to 20 for each group (Figure S3). To further study the contribution of *dsbA*, various organs were collected in order to determine if *E. faecalis* infection induced any strain-specific, pathological effects. To our surprise, cardiac microlesions were discovered within the myocardium that were devoid of immune cell infiltrate regardless of which *E. faecalis* strain was used (Figures 2A, B and S4). The microlesions appeared to form within roughly the same time frame (~24 h.p.i.) as cardiac microlesions that have previously been reported to occur during invasive infection with *S. pneumoniae* [11].

**Figure 2.**
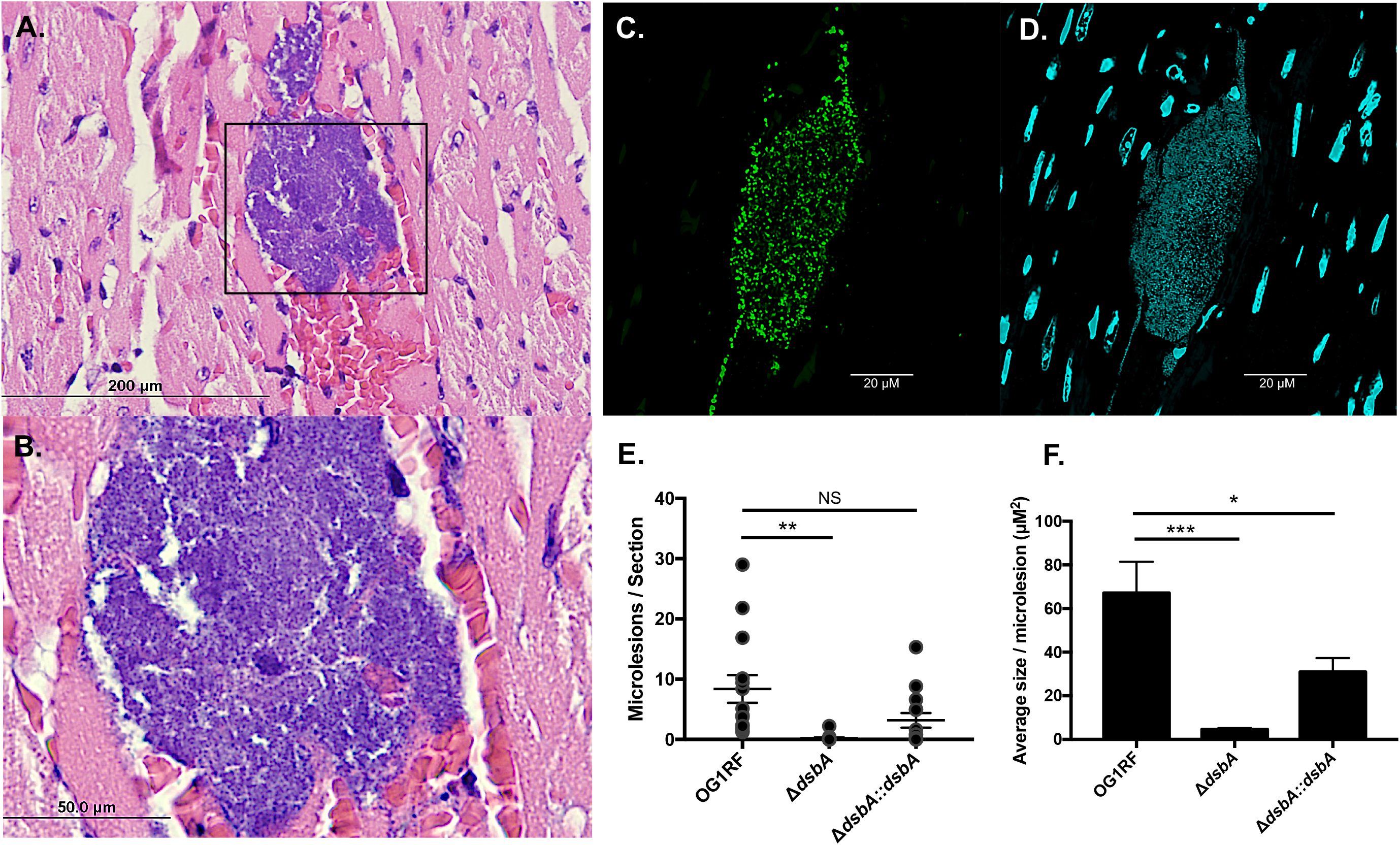
Cardiac microlesions form during invasive *E. faecalis* infection and are attenuated by loss of *dsbA*. Cardiac microlesions were observed using light microscopy of hematoxylin and eosin (H&E) stained paraffin embedded cardiac sections at 40X magnification **(A)**, and 100X magnification **(B)**. Immunofluorescent detection of wild-type *E. faecalis* was imaged using a 20X objective at 3X zoom (total of 60X magnification), a FITC conjugated anti-enterococcal antibody **(C)**, and DAPI for DNA staining **(D)**. Total microlesion counts **(E)**, and size measurements **(F)**, were obtained by quantification using light microscopy of H&E stained paraffin embedded cardiac sections from mice infected for 24-48 hours with *E. faecalis* (OG1RF n=14, Δ*dsbA* n=14, Δ*dsbA*::*dsbA* n=14). The data were averaged from 3 independent experiments and analyzed using *Student’s* two-tailed t-test (* indicate a *P* < 0.05, ** *P* < 0.01, *** a *P* < 0.001)

To verify that the observed microlesions were the result of infection by *E. faecalis*, immunofluorescent staining on paraffin embedded cardiac tissue sections from mice infected with *E. faecalis* with a polyclonal anti-enterococcal IgG antibody was performed and revealed the presence of dense aggregates of *E. faecalis* within the microlesions (Figure 2C). The presence of bacterial DNA was further verified by counterstaining with DAPI, revealing an accumulation of non-cardiomyocyte nuclear DNA within microlesions (Figure 2D). To determine the contribution of DsbA, cardiac microlesions found on hematoxylin and eosin stained paraffin embedded cardiac sections were quantified following infection with Δ*dsbA*. A significant attenuation in microlesion formation during infection with Δ*dsbA* was observed relative to wild-type, while no significant difference was observed in mice infected with the *dsbA* complement (Figure 2E). Furthermore, when the sizes of the cardiac microlesions were compared, the wild-type lesions were significantly larger than those found in the Δ*dsbA* infected mice (Figure 2F). Importantly, the size of the microlesions appeared to correlate with level of *dsbA* expression observed in Western blots of *E. faecalis* whole cell lysates (Figure 1B), with the Δ*dsbA*::*dsbA* strain manifesting an in-between phenotype to that of the "*dsbA* and wild-type strains, consistent with partial complementation. It is possible that less microlesion formation occurred with the Δ*dsbA* strain due poor growth in the blood, but we do not favor this model as in vitro growth curves in serum showed no defect (Figure S1B). In conclusion, these data demonstrate that cardiac microlesion formation occurs during systematic infection with *E. faecalis* and is dependent on DsbA.

### Apoptosis and necroptosis contribute to cardiomyocyte cell death within microlesions

In previous studies of pneumococcal cardiac microlesion formation, apoptosis was found to be associated with microlesion formation in mice [11]. A subsequent study determined that cardiomyocyte cell death occurs following exposure to pneumolysin (*ply*) and pneumococcal pyruvate oxidase (*spxB*) derived hydrogen peroxide [14]. Based on these observations and the fact that *E. faecalis* lacks *ply* and *spxB* homologs, we sought to determine whether *E. faecalis* could promote cell death of cardiomyocytes and identify the mechanism. To determine if apoptosis contributes to *E. faecalis* microlesion formation, we performed immunofluorescent microscopy on heart sections of infected mice to detect PARP [Poly(ADP-ribose) Polymerase], an enzyme that is proteolytically cleaved by the effector Caspase-3, and is one hallmark event of apoptosis. In agreement with previous reports, markers of apoptosis were observed to be concentrated at sites of microlesion formation (Figure 3A and S5).

**Figure 3.**
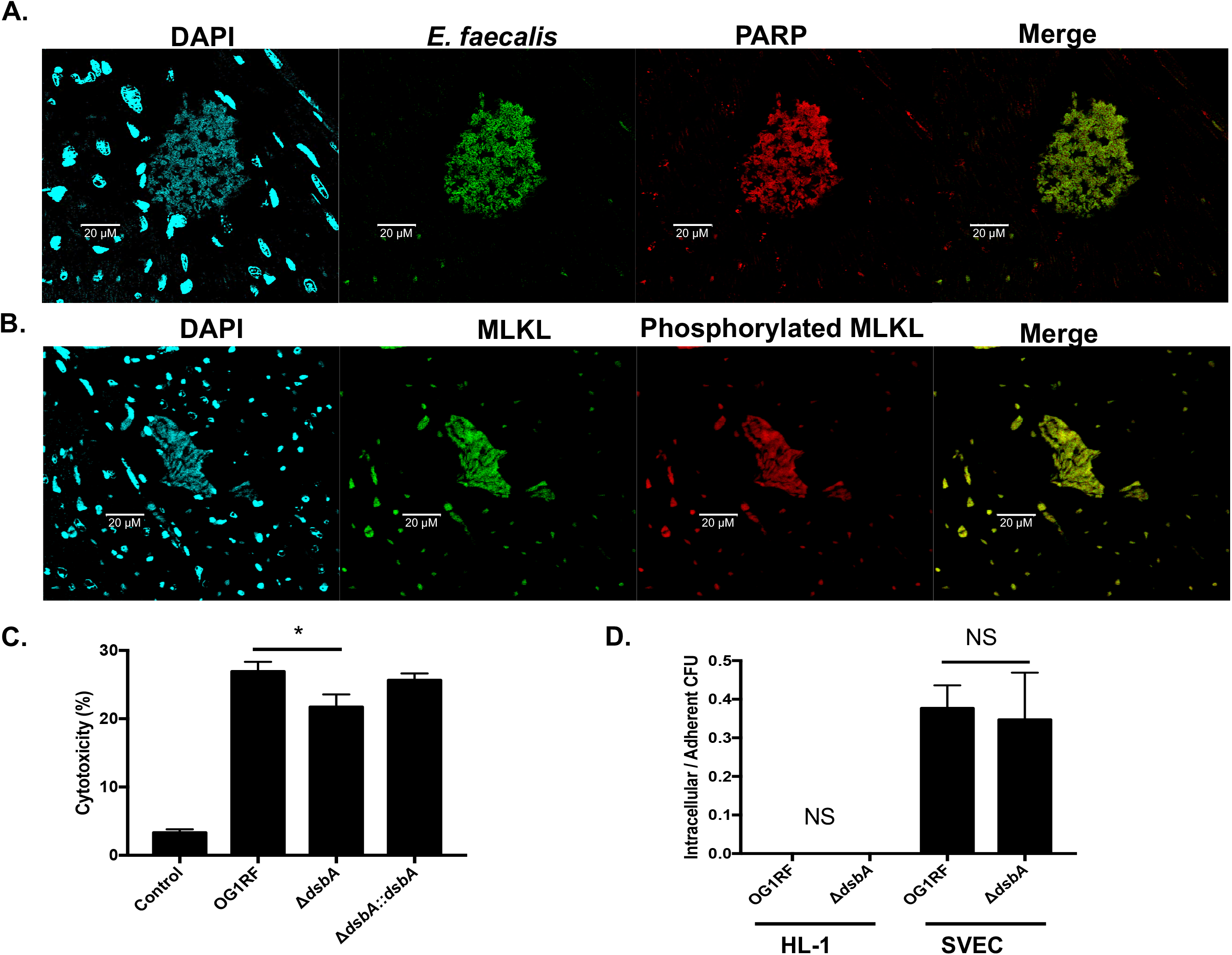
Necroptosis contributes to cardiotoxicity within cardiac microlesions. **(A)** Immunofluorescent microscopy of paraffin embedded cardiac cross sections at 20X were performed to detect *E. faecalis* using FITC conjugated anti-faecalis antibodies, and apoptosis using Alexa 594 conjugated anti-PARP antibodies at sites of microlesion formation, while DAPI was used to stain nuclei. **(B)** Immunofluorescent microscopy of paraffin embedded cardiac cross sections at 20X were performed to detect MLKL using FITC conjugated anti-MLKL antibodies, and Alexa 594 conjugated anti-phosphorylated MLKL antibodies at sites of microlesion formation, while DAPI was used to stain nuclei. **(C)** *E. faecalis* induced cell death in HL-1 cardiomyocytes 4 hours post-infection (h.p.i.) as determined by measuring lactate dehydrogenase. The data represent an average of 3 independent experiments and were analyzed using *Student’s* two-tailed t-test (* indicate a *P* < 0.05). **(D)** Comparison of OG1RF and Δ*dsbA* invasion rates into HL-1 cardiomyocytes and SVEC endothelial cells. The graph represents data from 3 independent experiments that were analyzed using *Student’s* two-tailed t-test

In addition to apoptosis, we hypothesized that necroptosis also contributes to *E. faecalis* mediated cardiac microlesion formation. Necroptosis is a programmed form of necrotic cell death that is generally inflammatory, and has been shown to contribute to the formation of *S. pneumoniae* microlesions within the myocardium of acutely ill mice and non-human primates [12]. Importantly, necroptosis is considered a key cell death pathway in cardiomyocytes during ischemia-reperfusion injury and acute coronary syndrome [23–25]. To test the hypothesis that necroptosis contributes to the formation of *E. faecalis* microlesions, we stained for markers of necroptosis [26], such as mixed lineage kinase domain-like pseudokinase (MLKL) and phosphorylated MLKL, on cardiac tissue sections obtained from infected mice. As a result, we detected concentrated signals within the microlesions (Fig 3B and Figure S6) that suggest that necroptosis does indeed contribute to *E. faecalis* cardiac microlesion formation.

To further look at the effects of *E. faecalis* and the *dsbA* mutant on heart cells, we infected HL-1 cardiomyocytes and subsequently measured lactate dehydrogenase (LDH) release into the cell culture supernatant to assess cell death. As shown in Figure 3C, cell death of the cardiomyocytes is significantly decreased following exposure to the *dsbA* mutant compared to wild-type suggesting that DsbA contributes to cardiac cell death (Figure 3C). While *E. faecalis* promoted death of the cardiomyocytes, antibiotic protection assays indicated that both the wild-type and *dsbA* mutant bacteria were unable to invade cardiomyocytes relative to endothelial cells (Figure 3D). Taken together, these results suggest that during bacteremia, *E. faecalis* is able to translocate across the vascular endothelium in order to enter the myocardial tissue. Once there, *E. faecalis* is unable to invade cardiomyocytes, but can form microcolonies that promote necroptosis in addition to apoptosis. Whether other cell death pathways additionally contribute will be fodder for future investigations.

### Cardiomyocyte inflammatory cytokine response is suppressed following exposure to *E. faecalis*

Cardiac microlesions are characterized by bacteria filled vacuoles that are largely devoid of immune cell infiltrate [11, 27]. Previous efforts to explain the lack of an immune response to pneumococcal microlesions have yielded interesting insights. For example, *S. pneumoniae* within microlesions are thought to subvert the host response through a biofilm-mediated mechanism involving the pneumolysin mediated killing of resident macrophages, thus creating a more permissive environment for pneumococcal growth and survival [13]. There are also examples of *E. faecalis* subverting the immune response. In previous work, it was shown that *E. faecalis* can survive within mouse peritoneal macrophages for extended periods of time [28] and can also resist phagosome acidification and autophagy [29]. Additionally, *E. faecalis* was shown to activate Phosphatidylinositol 3-Kinase signaling that inhibits apoptosis [30]. Because *E. faecalis* does not encode a pneumolysin homolog, nor does it secrete toxins known to kill macrophages or other immune cells, we wanted to examine the immune response to *E. faecalis* within cardiomyocytes. Therefore, following infection with *E. faecalis*, inflammatory chemokine and cytokine levels were measured. Wild-type *E. faecalis* was discovered to promote an immune quiescent response, while the *dsbA* mutant caused a more inflammatory immune response (Figure 4A and Table S2). Specifically, infection of cardiomyocytes with wild-type increased the expression of two pro-inflammatory cytokines IL-4 and b-NGF cytokines [31, 32]. Modest increases were also observed in VEGF, an important determinant of sepsis morbidity and mortality [33], and GM-CSF that serves as an immunomodulatory cytokine [34]. In contrast, cardiomyocytes infected with Δ*dsbA* led to increases in Rantes, IFNr, MIP-1a, and IL-12, with modest increases also observed in Leptin and G-CSF. With the exception of G-CSF, an immunomodulatory cytokine with roles in cardiac cell generation and repair [35, 36], all of the other aforementioned cytokines are pro-inflammatory [37–40]. These results implicate the contribution of specific host signaling pathways that are normally suppressed in the presence of DsbA during infection, as modeled in Figure 4B. Collectively, these data suggest that *E. faecalis* is capable of suppressing the host cardiomyocyte immune response.

**Figure 4.**
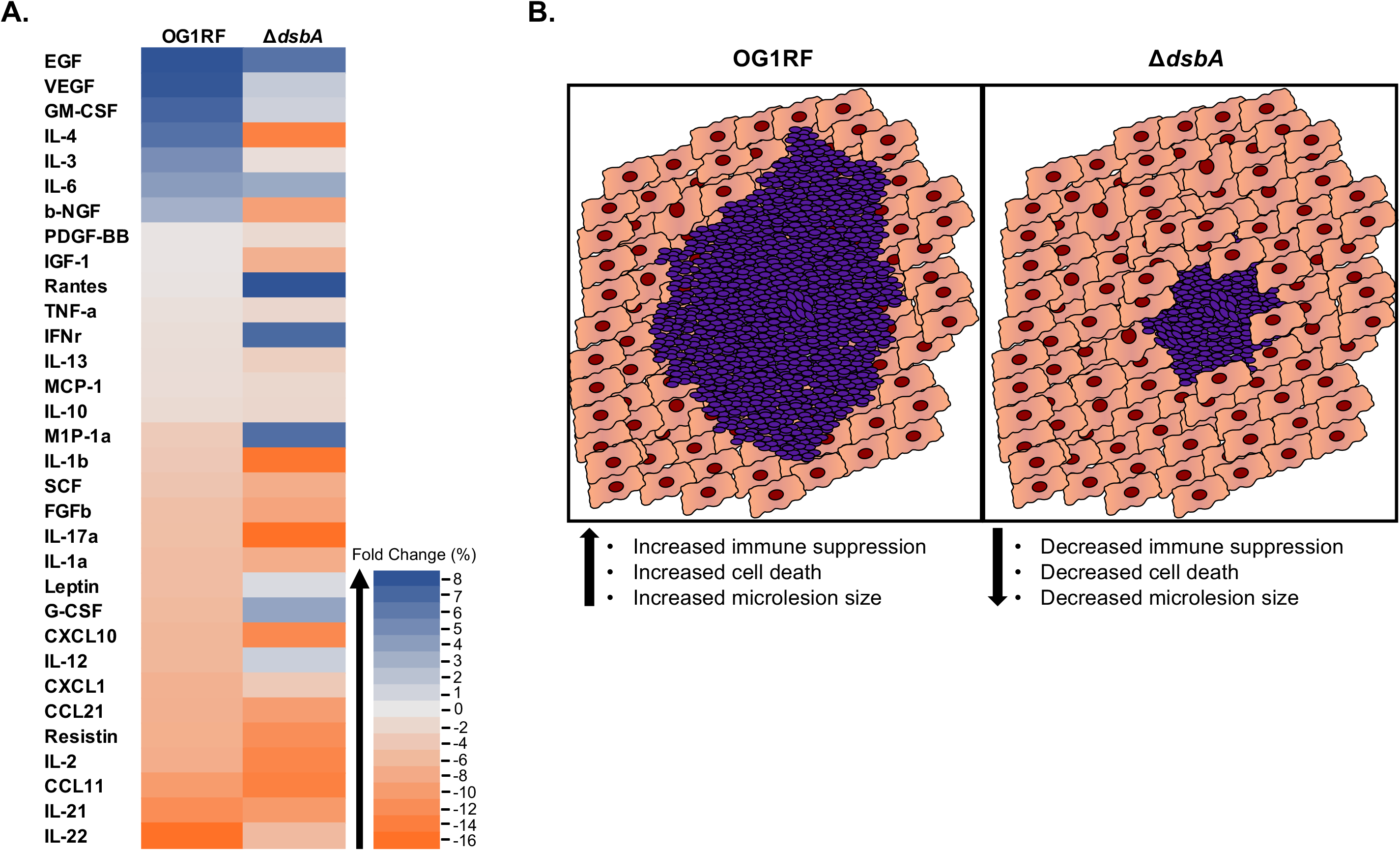
Wild-type *E. faecalis* suppresses the cardiomyocyte inflammatory response *in-vitro*. **(A)** HL-1 cardiomyocytes exposed to *E. faecalis* promote a suppressed inflammatory response while Δ*dsbA* infected cardiomyocytes yield a more inflammatory response. Data was obtained from the fold-expression change analysis (relative to uninfected controls) of at least 3 independent experiments for each strain. **(B)** Model of *E. faecalis* mediated suppression of the host cardiomyocyte immune response and effect on microlesion size. Wild-type *E. faecalis* is postulated to downregulate the cardiomyocyte immune response more effectively than the *dsbA* mutant resulting in more cell death and increased lesion size. However, both strains cause cell death by apoptosis and necroptosis and inhibit immune cell infiltration

## Discussion

In this work, we demonstrated for the first time that cardiac microlesions form during severe *E. faecalis* bacteremic infection of mice. Importantly, microlesions occurred despite *E. faecalis* not encoding homologs of the pneumococcal virulence determinants implicated in microlesion formation. These data suggest that *E. faecalis* may be expressing functionally redundant proteins that serve to promote similar interactions to *S. pneumoniae* in mediating microlesion formation, such as CbpA and pneumolysin, which facilitate vascular endothelial cell translocation and cardiomyocyte cell death, respectively. Indeed, while *E. faecalis* does not encode a CbpA homolog, it does encode other proteins capable of binding host extracellular matrix proteins. These include EfbA, which binds fibronectin [41], ACE, which binds laminin receptor [42], and Ebp pili that bind fibrinogen [43].

Interestingly, we did discover one factor, DsbA, that contributes to microlesion formation and inhibits the cardiomyocyte inflammatory immune response, as modeled in Figure 4B. Dsb proteins work to catalyze disulfide bond formation in proteins and have been best studied in Gram-negative bacteria [44, 45]. While Dsb proteins have established importance in the biology of certain Gram-positive bacteria, Dsb proteins are not well studied in Firmicutes like *E. faecalis* and have generally not been considered important [44]. Most of these species secrete few proteins containing cysteines and some, notably *S. pneumoniae*, do not encode any thiol-disulfide oxidoreductases [46, 47]. Thus, it will be of interest to further investigate the role of Dsb proteins in these species.

Our prior work established EntV as a DsbA substrate, currently the only one identified [21]. However, loss of EntV did not reduce pathogenicity in the worm model (Figure 1A), and exposure to EntV, even at very high concentrations, was non-toxic to mammalian cells [22]. As such, we do not favor EntV as being the source of DsbA’s effects. While the specific *E. faecalis* proteins involved in modulating the host immune response are unknown, we speculate that the Fibronectin-binding protein (Efb) and Choline binding protein may contribute to cardiac microlesion formation by serving as functionally redundant proteins to pneumococcal CbpA, a virulence determinant known to be integral to microlesion formation [11]. Moreover, both proteins contain 5 cysteines and are expressed extracellularly. Thus, our future research efforts will be geared toward identifying additional substrates of DsbA and to determining their contribution to microlesion formation.

Pneumococcal ChoP interactions with host PAFr were determined to be a prerequisite for vascular endothelial cell invasion during IPD. In light of our observations of *E. faecalis* invasion of vascular endothelial cells despite an absence of this wall teichoic acid structure, it is possible that another host receptor is required for endothelial cell invasion. Interestingly, recent studies focused on bacterial cell wall entry and signaling in eukaryotic cells have identified a PAFr-independent mechanism of tissue invasion that is actin-dependent and engenders Rac1, Cdc42, and phosphatidylinositol 3-kinase (PI3K) signaling leading to cell wall internalization via a macropinoctytosis-like pathway [48]. Whether such a pathway contributes to microlesion formation is unknown. In the future, it will be of interest to study the mechanisms of tissue invasion and the host signaling pathways involved.

Although the role of cardiac microlesion formation in humans during invasive disease remains unclear, cardiac microlesion formation is becoming increasingly appreciated as a risk factor [49]. To our knowledge, no one has investigated whether cardiac microlesions form in humans during enterococcal infection. We speculate that formation occurs and contributes to cardiac abnormalities or death in patients with enterococcal infective endocarditis or sepsis [1, 3–9]. This study builds on our understanding of cardiac microlesions by establishing *E. faecalis* as another etiological agent capable of causing them in a vertebrate animal model. Considering that *E. faecalis* microlesion pathology appears identical to that caused by *S. pneumoniae*, it is likely that scar formation occurs following resolution of infection, potentially predisposing individuals to future adverse cardiac events [11]. Taken together, these data highlight the need for further investigation in order to understand the impact of microlesion formation on the heart and mortality during invasive *E. faecalis* infection.

---

**Table S1.** Strains used in this study.

| *Enterococcus faecalis* Strains |  |  |  |  |
| --- | --- | --- | --- | --- |
| Strain | Genotype | Characteristics | Parent | Source |
| OG1RF | Wild-type | Gel<sup>+</sup> Spr<sup>+</sup> Rf<sup>R</sup> Fa<sup>R</sup> | OG1 | [50] |
| AOBD1 | $\Delta dsbA$ | Gel<sup>+</sup> Spr<sup>+</sup> Rf<sup>R</sup> Fa<sup>R</sup> | OG1 | [21] |
| AOBD2 | $\Delta dsbA::dsbA$ | Gel<sup>+</sup> Spr<sup>+</sup> Rf<sup>R</sup> Fa<sup>R</sup> | OG1 | [21] |

**Table S2.** Cytokine Profiling Expression Data.

|  | Exp. 1 | Exp. 2 | Exp. 3 |  |  |
| --- | --- | --- | --- | --- | --- |
|  | OG1RF | OG1RF | OG1RF | Average | Std. Dev |
| TNF-a | -7% | 7% | -2% | -1% | 7% |
| IGF-1 | 2% | -7% | 4% | 0% | 6% |
| VEGF | 23% | 4% | -2% | 8% | 13% |
| IL-6 | 7% | 5% | 1% | 4% | 3% |
| FGFb | 11% | -28% | 0% | -5% | 20% |
| IFNr | 5% | 3% | -11% | -1% | 9% |
| EGF | -10% | 33% | 2% | 8% | 23% |
| Leptin | 0% | -16% | -1% | -6% | 9% |
| IL-1a | -7% | -14% | 4% | -6% | 9% |
| IL-1b | -8% | -22% | 18% | -4% | 20% |
| G-CSF | 1% | -22% | 2% | -6% | 14% |
| GM-CSF | -10% | 19% | 14% | 7% | 16% |
| MCP-1 | -7% | -8% | 11% | -1% | 11% |
| M1P-1a | 3% | 3% | -17% | -4% | 12% |
| SCF | -15% | 5% | -3% | -5% | 10% |
| Rantes | -13% | 19% | -7% | 0% | 17% |
| PDGF-BB | -16% | 8% | 7% | 0% | 13% |
| b-NGF | -20% | 12% | 17% | 3% | 20% |
| IL-17a | -22% | -4% | 9% | -5% | 16% |
| IL-2 | -9% | -10% | -5% | -8% | 3% |
| IL-4 | 12% | -1% | 9% | 7% | 7% |
| IL-10 | 11% | -13% | -3% | -2% | 12% |
| Resistin | -13% | -3% | -6% | -8% | 5% |
| IL-12 | -3% | 0% | -17% | -7% | 9% |
| CCL11 | -9% | -25% | 3% | -10% | 14% |
| CCL21 | -17% | 0% | -5% | -7% | 9% |
| IL-3 | 8% | 3% | 5% | 5% | 2% |
| IL-13 | -7% | -7% | 10% | -1% | 10% |
| IL-21 | -24% | -14% | 1% | -12% | 13% |
| IL-22 | -18% | -19% | -12% | -16% | 4% |
| CXCL10 | -12% | -10% | 2% | -7% | 8% |
| CXCL1 | -5% | -4% | -13% | -7% | 5% |

\*OG1RF ordered from Largest to smallest

| | OG1RF | $\Delta dsbA$ |
| --- | --- | --- |
| EGF | 8% | 3% |
| VEGF | 8% | 1% |
| GM-CSF | 7% | 1% |
| IL-4 | 7% | -12% |
| IL-3 | 5% | -1% |
| IL-6 | 4% | 2% |
| b-NGF | 3% | -8% |
| PDGF-BB | 0% | -2% |
| IGF-1 | 0% | -6% |
| Rantes | 0% | 4% |
| TNF-a | -1% | -2% |
| IFNr | -1% | 4% |
| IL-13 | -1% | -3% |
| MCP-1 | -1% | -2% |
| IL-10 | -2% | -2% |
| M1P-1a | -4% | 3% |
| IL-1b | -4% | -13% |
| SCF | -5% | -7% |
| FGFb | -5% | -8% |
| IL-17a | -5% | -14% |
| IL-1a | -6% | -7% |
| Leptin | -6% | 0% |
| G-CSF | -6% | 2% |
| CXCL10 | -7% | -11% |
| IL-12 | -7% | 1% |
| CXCL1 | -7% | -3% |
| CCL21 | -7% | -9% |
| Resistin | -8% | -10% |
| IL-2 | -8% | -11% |
| CCL11 | -10% | -12% |
| IL-21 | -12% | -9% |
| IL-22 | -16% | -5% |

|  | Exp. 1 | Exp. 2 | Exp. 3 |  |  |
| --- | --- | --- | --- | --- | --- |
| | $\Delta dsbA$ | $\Delta dsbA$ | $\Delta dsbA$ | Average | Std. Dev |
| TNF-a | -2% | 5% | -8% | -2% | 6% |
| IGF-1 | -15% | -7% | 3% | -6% | 9% |
| VEGF | 2% | 1% | 0% | 1% | 1% |
| IL-6 | 0% | 13% | -8% | 2% | 10% |
| FGFb | -7% | -15% | -2% | -8% | 7% |
| IFNr | 16% | 7% | -12% | 4% | 14% |
| EGF | -8% | 31% | -12% | 3% | 24% |
| Leptin | 13% | -8% | -4% | 0% | 11% |
| IL-1a | -17% | -10% | 7% | -7% | 12% |
| IL-1b | -20% | -19% | -1% | -13% | 11% |
| G-CSF | -12% | 14% | 3% | 2% | 13% |
| GM-CSF | -18% | 18% | 2% | 1% | 18% |
| MCP-1 | -20% | -9% | 23% | -2% | 22% |
| M1P-1a | -6% | 3% | 13% | 3% | 10% |
| SCF | -10% | -20% | 10% | -7% | 15% |
| Rantes | -3% | 20% | -4% | 4% | 14% |
| PDGF-BB | -23% | 15% | 4% | -2% | 20% |
| b-NGF | -16% | -10% | 1% | -8% | 9% |
| IL-17a | -45% | -1% | 4% | -14% | 27% |
| IL-2 | -35% | -1% | 1% | -11% | 20% |
| IL-4 | -17% | -21% | 2% | -12% | 13% |
| IL-10 | 0% | -2% | -4% | -2% | 2% |
| Resistin | -19% | -6% | -6% | -10% | 7% |
| IL-12 | -13% | 14% | 1% | 1% | 14% |
| CCL11 | -13% | -30% | 7% | -12% | 18% |
| CCL21 | -33% | 4% | 3% | -9% | 21% |
| IL-3 | -8% | 2% | 3% | -1% | 6% |
| IL-13 | -25% | -3% | 19% | -3% | 22% |
| IL-21 | -23% | -10% | 6% | -9% | 15% |
| IL-22 | -26% | -7% | 18% | -5% | 22% |
| CXCL10 | -14% | -17% | -2% | -11% | 8% |
| CXCL1 | -12% | -15% | 17% | -3% | 17% |

**Supplemental Figure 1. *E. faecalis* growth curve. (A)** *E. faecalis* strains were grown for 15 hours in BHI and absorbance (OD_600nm_) measured every 10 minutes for 16 hours. **(B)** *E. faecalis* grown in serum and absorbance (OD_600nm_) measured every 10 minutes for 16 hours. Figures are representative of 3 independent experiments.

**Supplemental Figure 2. Coomassie stained SDS-PAGE gel of *E. faecalis* whole cell lysates separated by electrophoresis. (A)** SDS-PAGE separation of *E. faecalis* whole cell lysates was stained with Coomassie to verify that each well received 10 µg / lane of whole cell lysate. Coomassie stained gel is the loading control for the Western blot in Figure 1B.

**Supplemental Figure 3. DsbA is required for *E. faecalis* pathogenesis. (A)** Mice (n=17-20) challenged with 3.8 − 4.2 × 10^8^ CFUs of *E. faecalis* in a mouse model of peritonitis were assessed for survival over 96 hours. Mice infected with Δ*dsbA E. faecalis* lived significantly longer (*P* = 0.0051) than mice challenged with wild-type or the complemented strain (no significant difference). Survival curve was calculated from 3 independent experiments using log-rank analysis.

**Supplemental Figure 4. (A)** 40X magnification image of cardiac microlesion formation in mice infected with Δ*dsbA.* **(B)** 100X image of cardiac microlesion in Figure S4A. **(C)** 40X magnification image of cardiac microlesion formation in mice infected with Δ*dsbA*::*dsbA*. **(D)** 100X image of cardiac microlesion in Figure S4C.

**Supplemental Figure 5. Necroptosis contributes to cardiotoxicity within cardiac microlesions. (A)** Immunofluorescent microscopy of paraffin embedded cardiac cross sections using a 20X objective at 4X Zoom (80X total magnification) were performed to detect MLKL using FITC conjugated anti-MLKL antibodies, and Alexa 594 conjugated anti-phosphorylated MLKL antibodies at sites of microlesion formation, while DAPI was used to stain nuclei.

**Supplemental Figure 6. Apoptosis contributes to cell death within cardiac microlesions. (A)** Immunofluorescent microscopy of paraffin embedded cardiac cross sections using a 20X objective at 4X Zoom (80X total magnification) were performed to detect *E. faecalis* using FITC conjugated anti-*E. faecalis* antibodies, and Alexa 594 conjugated anti-PARP antibodies at sites of microlesion formation, while DAPI was used to stain nuclei.

## Meetings

This work has not been presented.

## Corresponding author contact information

Danielle A. Garsin, Ph.D.

6431 Fannin St.

MSB 1.168

Houston, TX 77030

Telephone: (713) 500-5454

Fax: (713) 500-5499

