## Supplemental Figures for "Cardiac Microlesions Form During Severe Bacteremic *Enterococcus faecalis* Infection"

Supplemental Figure 1

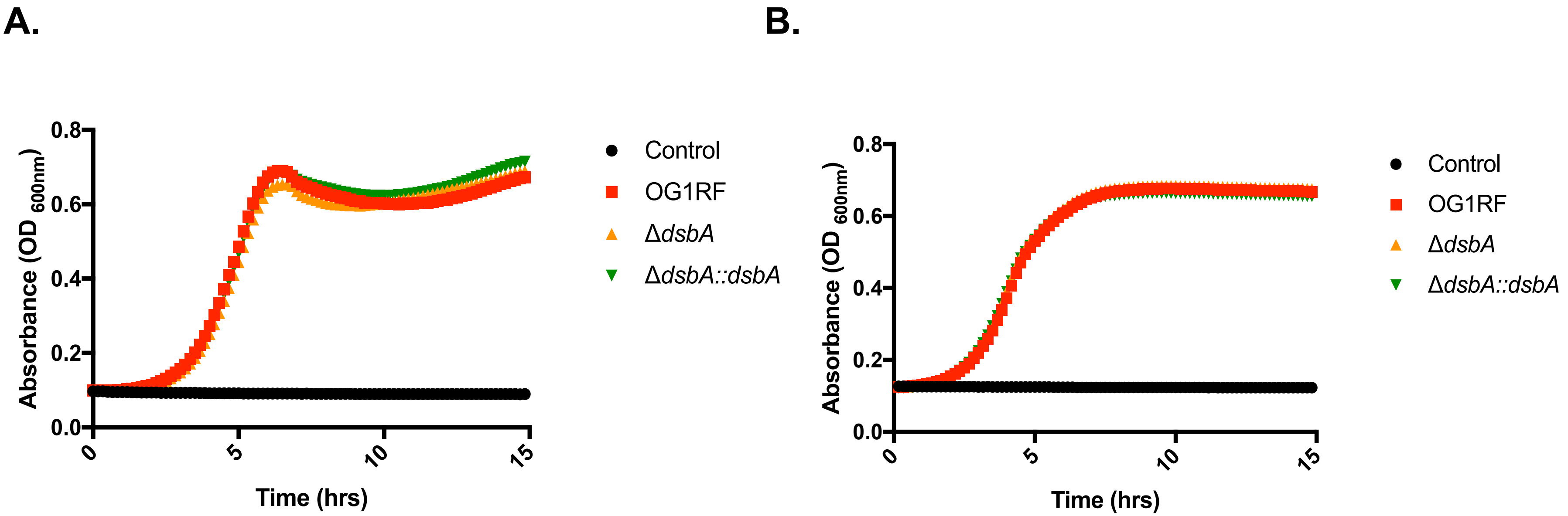

Supplemental Figure 2

A.

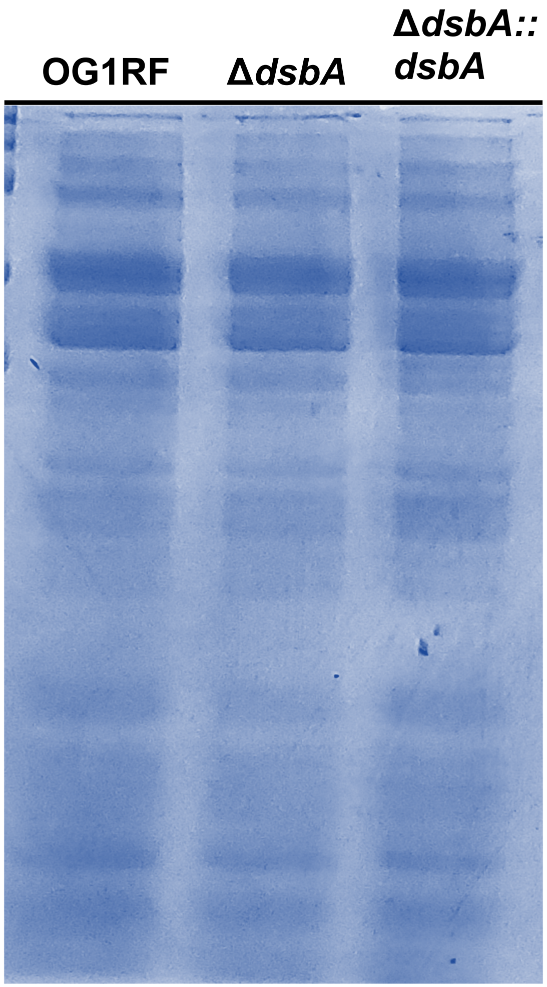

Supplemental Figure 3

**A.**

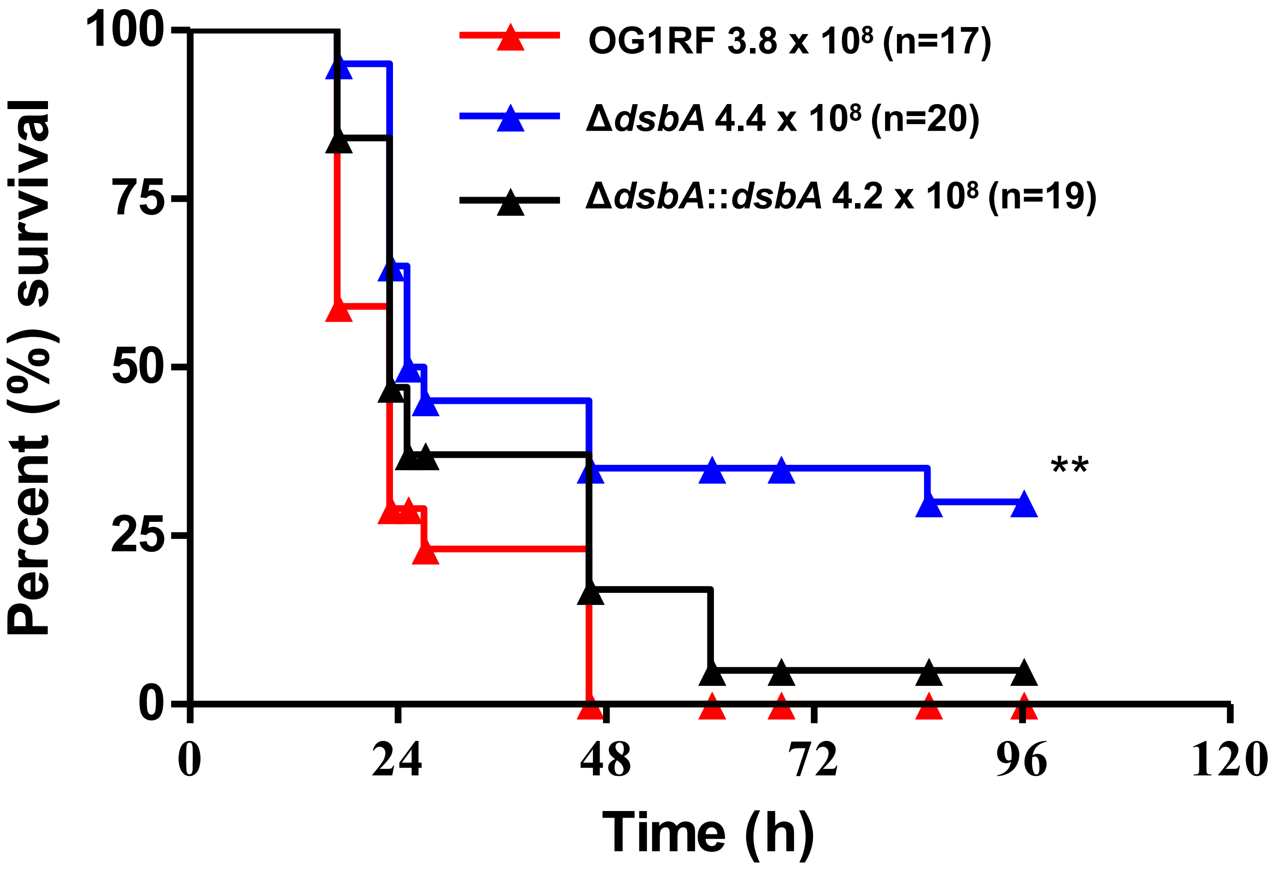


**OG1RF vs Δ*dsbA* = *P* 0.0051**

**Δ*dsbA* vs Δ*dsbA*::*dsbA* = *P* 0.0715**

Supplemental Figure 4

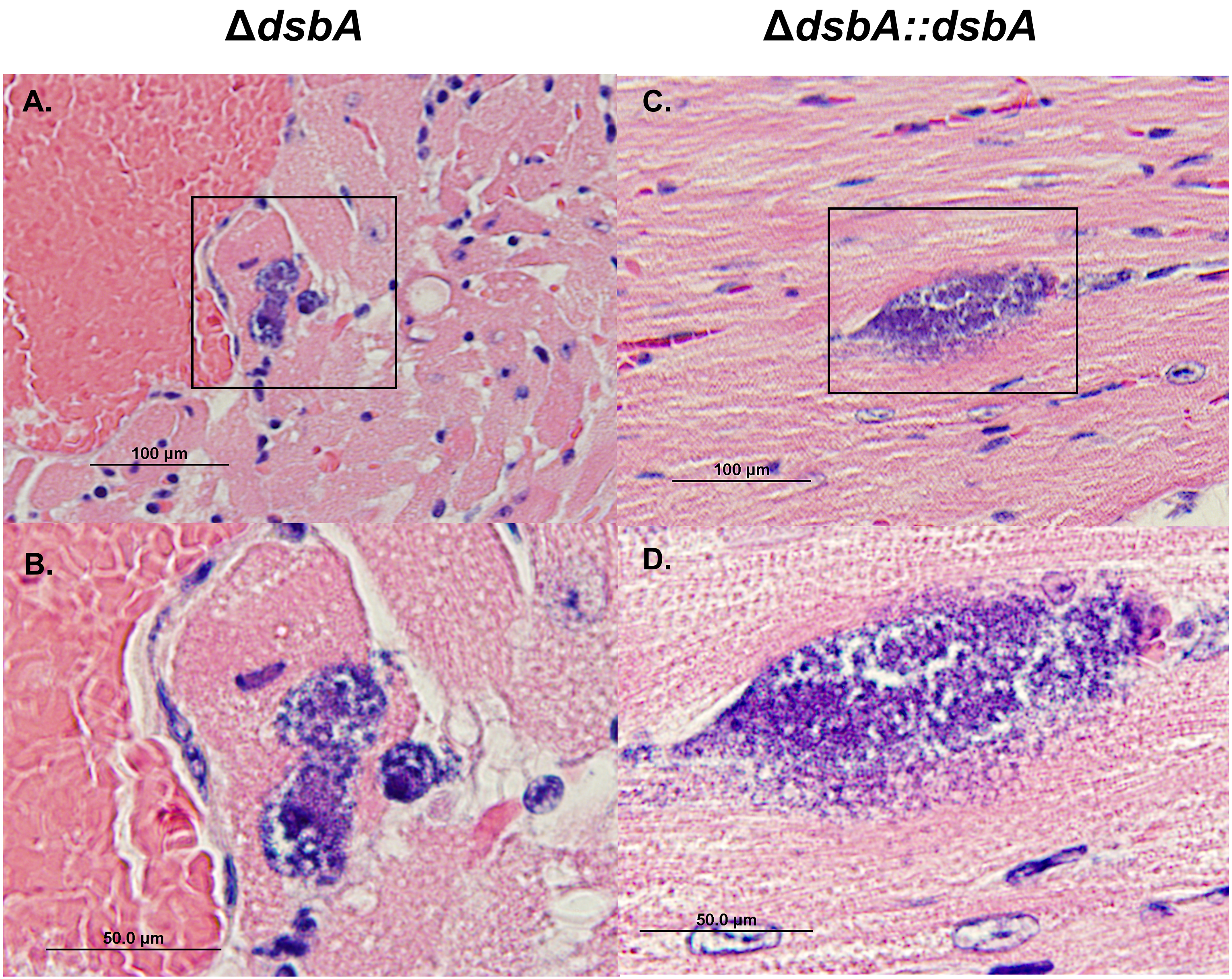

Supplemental Figure 5

**A.**

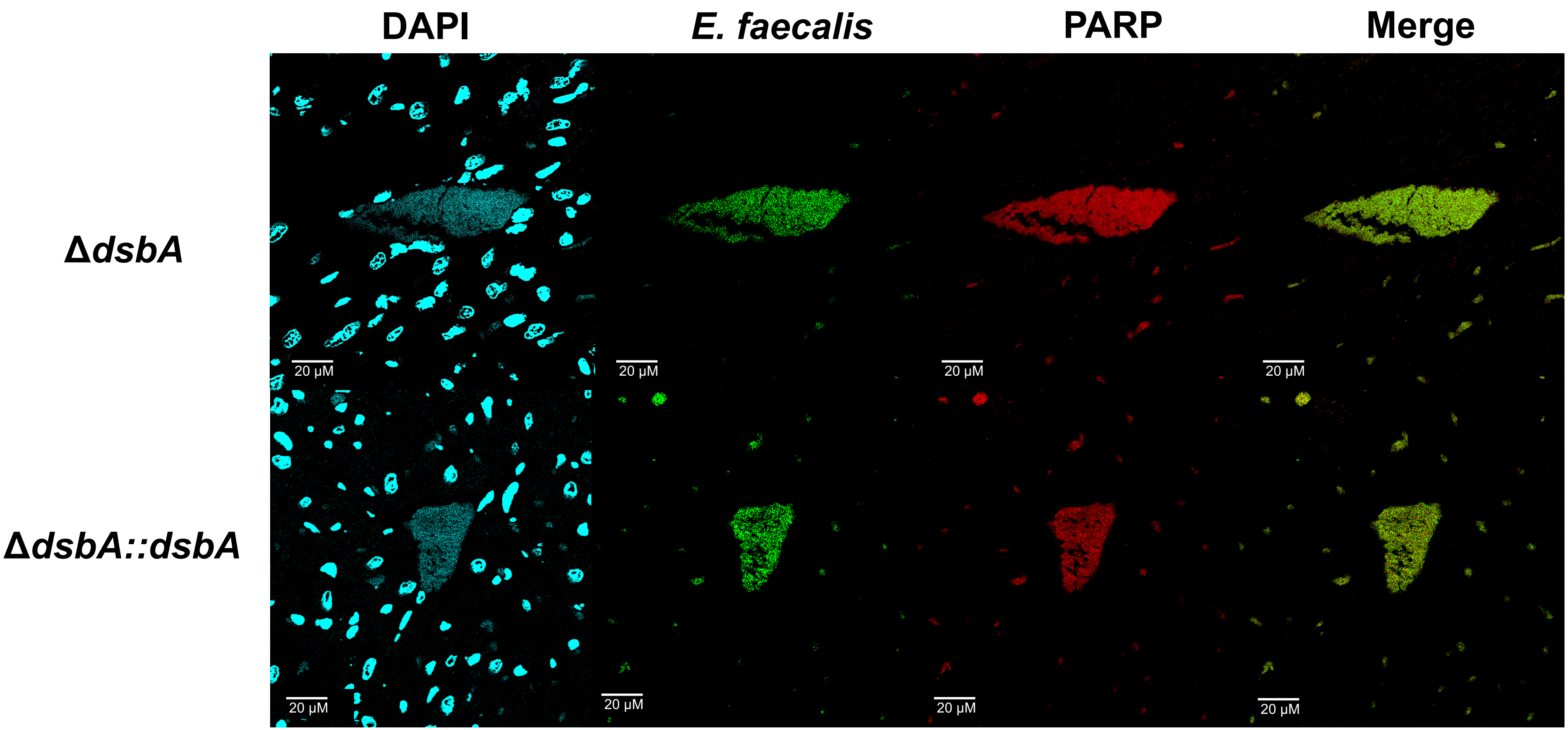

Supplemental Figure 6

**A.**

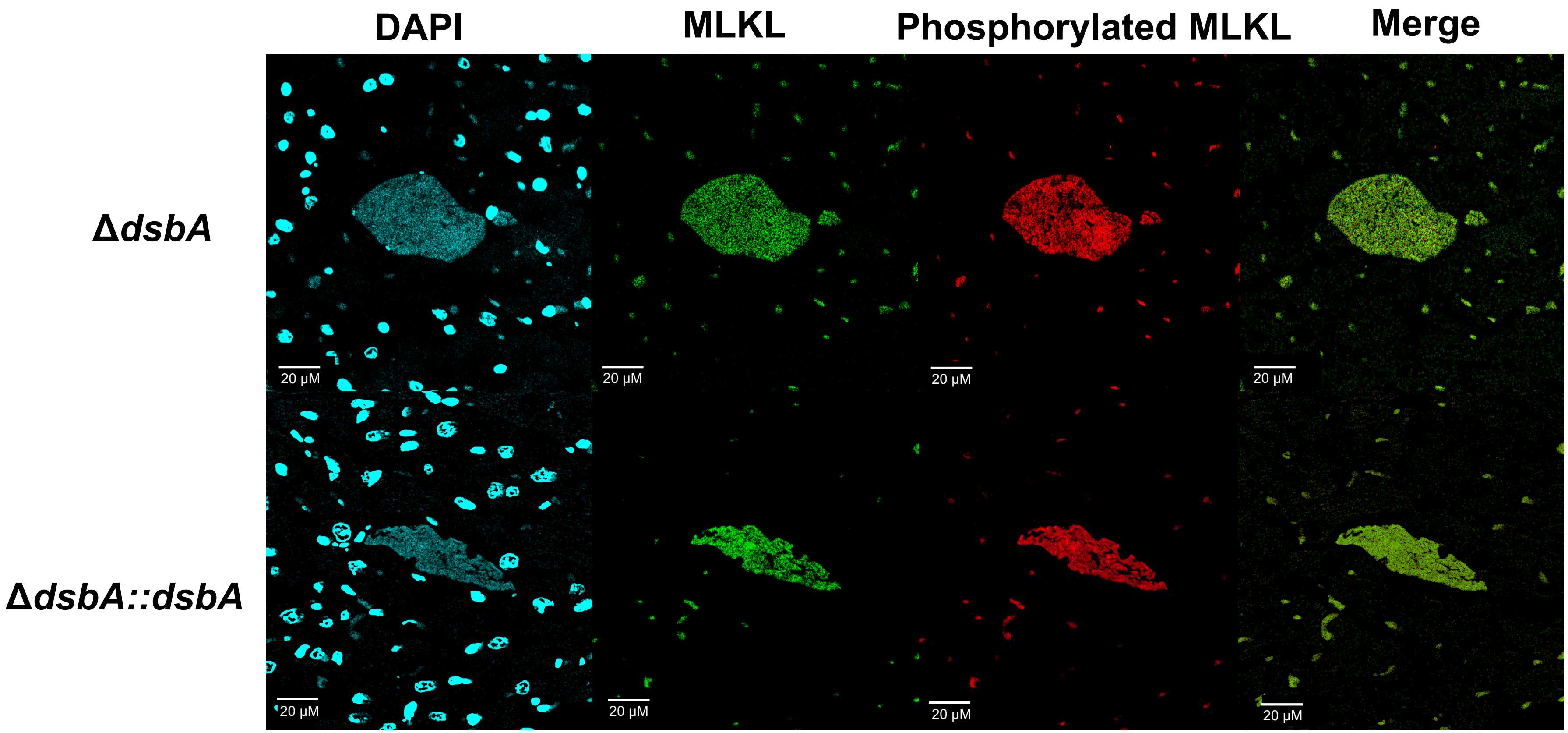
